# Phosphorylation regulates auto-inhibition of kinesin KIF3A

**DOI:** 10.1101/503680

**Authors:** Kunning Chen, Wuan-Geok Saw, Dilraj Lama, Chandra Verma, Gerhard Grüber, Cheng-Gee Koh

## Abstract

Kinesin are molecular motors that move along the microtubules. They function to transport cargoes, vesicles and organelles to designated locations in the cells. KIF3A belongs to the Kinesin-2 family and forms a heterotrimeric complex with KIF3B and KAP3. We have earlier shown that the cargo trafficking activity of KIF3 motor can be regulated by CaMKII kinase and POPX2 phosphatase. In this study, we elucidated the mechanism of KIF3A regulation. We find that KIF3A adopts an auto-inhibited state through the interaction between the motor and tail domains. The motor-tail interaction also hinders the ATPase activity of the motor domain. We show that the phosphorylation status of serine-689/690 (mouse/human) at the C-terminal region of KIF3A is crucial for the motor-tail interaction. The motor domain does not interact well with the tail domain when serine-689 is phosphorylated by CaMKII or mutated to aspartic acid to mimic phosphorylation. Molecular dynamics simulations suggest that the non-phosphorylated tail domain folds into the hydrophobic pocket formed by the motor dimers. Phosphorylation of serine-689 results in conformational changes that leads to the relieve of auto-inhibition.

## INTRODUCTION

Intracellular transport is an important process to designate large molecules, vesicles and organelles to the correct cellular locations where they are needed. Kinesins are molecular motors that enable intracellular transport along the microtubules (1). To date, 45 kinesin superfamily members have been identified and classified into 15 families. Kinesin family members share similar characteristics. They are composed of a motor domain which binds and moves along microtubule tracks with energy released from ATP hydrolysis, a tail domain that binds to cargoes, and a coiled-coil stalk region that links the two domains (2). KIF3 is a family member of the Kinesin-2 family. Unlike the homodimeric KIF17 (3), which is another Kinesin-2 member, KIF3 forms a heterotrimeric complex. The KIF3 heterotrimeric complex is comprised of two motor subunits—KIF3A and KIF3B and a non-motor subunit, kinesin-associated protein KAP3 (4). KIF3C has also been reported to be a subunit of the KIF3 motor complex. KIF3C associates with KIF3A, but not KIF3B, and functions both in complex with KIF3A or independently from the KIF3 motor complex (5). Several cancer-related proteins such as the PAR-3 polarity complex, N-cadherin, β-catenin and adenomatous polyposis coli (APC) have been identified as cargoes of the KIF3 motor (6,7). KIF3 motor complex is known to be involved in intra-flagellar transport, cilium assembly, and cargo transportation in neurons. Loss of KIF3 leads to early developmental defects (8-12).

The activities of several kinesin families are reported to be regulated by autoinibition (13-16). In an autoinhibited state, kinesin forms a folded, compact conformation that cannot hydrolyze ATP or associate with cargos or microtubules. When activated, kinesin is released from autoinhibition and transformed into an active, extended configuration. Autoinhibition can be relieved by phosphorylation. Phosphorylation of threonine-937 on KIF11 by CDK1 (cyclin-dependent kinase 1) activates the motor and increases its microtubule-binding efficiency (17). TTK (dual specificity protein kinase) or CDK1-cyclin B phosphorylates CENP-E (centrosome-associated protein E) and causes it to unfold into an extended conformation and increases its processive motility along microtubules (18). The function of KIF3A motor subunit is also regulated by phosphorylation. CaMKII (Ca^2+^/calmodulin-dependent protein kinase) and POPX2 (Partner of PIX 2) phosphatase have been identified as the kinase and phosphatase pair that is responsible for the regulation of phosphorylation at serine-690 of KIF3A (19). KIF3A’s cargo transportation is found to be inhibited when serine-690 is mutated to alanine (19). In addition, it has been reported that the KIF3 complex possibly adopts both folded and extended conformations (20). Based on these findings, we propose that the KIF3A motor is autoinhibited when serine-690/serine-689 located at the C-terminal region of KIF3A (human/mouse) is in a dephosphorylated state and phosphorylation can relieve this autoinhibition.

## RESULTS

### Residue serine-689 residue of KIF3A (mouse) is phosphorylated by CaMKII

The residue serine-690 at the C-terminal tail of KIF3A was identified as a major phosphorylation site by various studies (21-25). CaMKII has been reported to be the kinase that phosphorylates KIF3A *(Homo sapiens)* at serine-690 (19). However, *Ichinose et al*. showed in their LC-MS/MS results that threonine-694 and serine-698 of the *Mus musculus* KIF3A were the phosphorylation sites for CaMKII, while serine-689 (equivalent to human serine-690) is phosphorylated by PKA (26). Serine-689 (mouse) or serine-690 (human) is among the conserved amino acid sequence RKRS at the C-terminal tail domain of KIF3A. This conserved sequence fits the consensus recognition sequence for CaMKII (RXXS/T) while neither threonine-694 nor serine-698 falls in the recognition motif of CaMKII (Fig. 1A).

**Figure 1.**
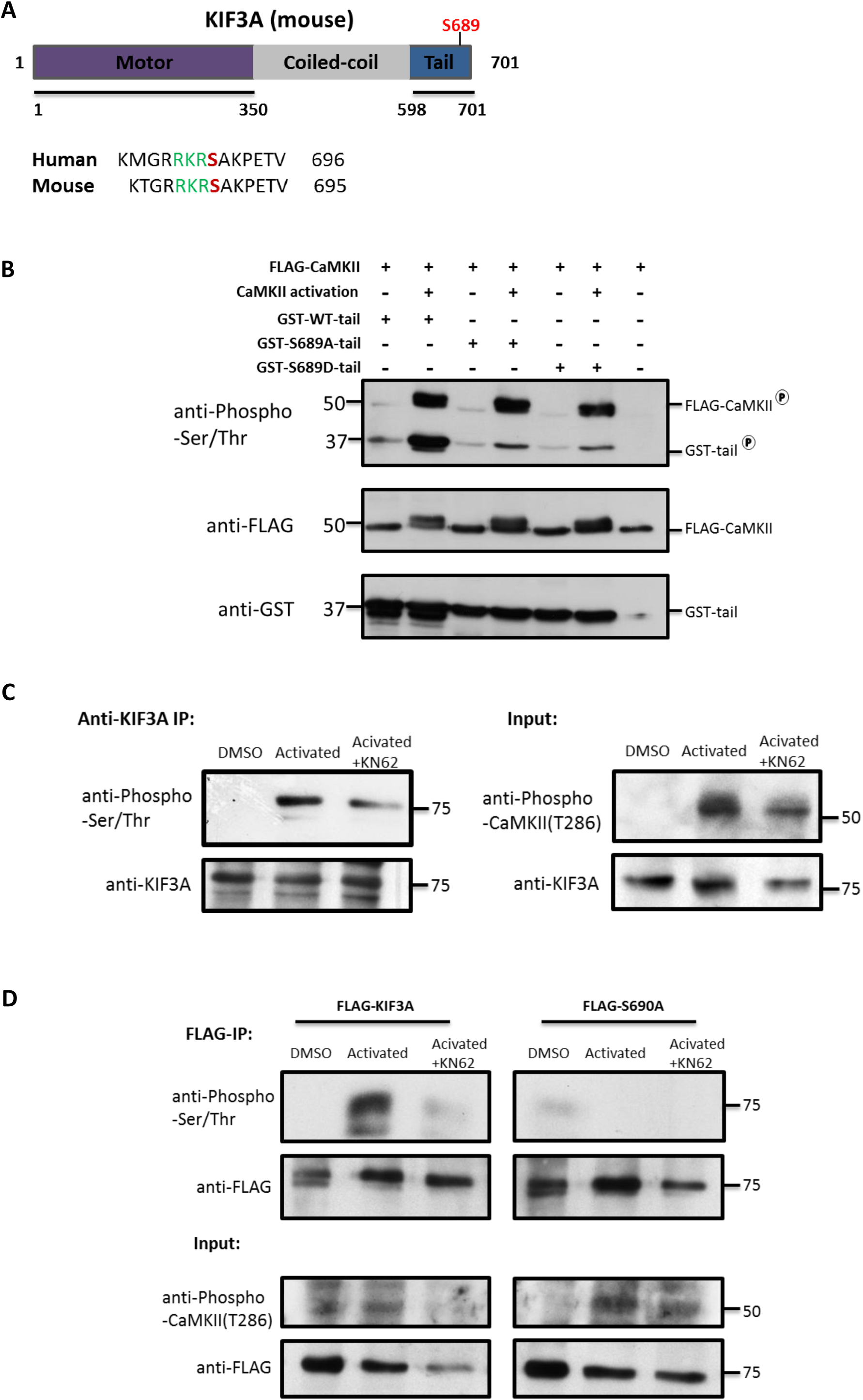
CaMKII specifically phosphorylates KIF3A on serine-689. (A) Domain organization of the mouse KIF3A. Mouse and human KIF3A show high sequence homology. Serine-689 of mouse KIF3A is equivalent to serine-690 of the human KIF3A. The conserved CaMKII recognition sequence RXXS/T is marked by green characters and the conserved serine residue is highlighted in red in both human and mouse sequences. (B) Activated CaMKII phosphorylates KIF3A tail domain. Inactive FLAG-CaMKII or active FLAG-CaMKII were incubated with 5 μg of GST-WT-tail, GST-S689A-tail or GST-S689D-tail respectively for 45 min at 30 °C. The reaction mixes were separated by SDS-PAGE and subjected to Western analysis with the respective antibodies. (C) For CaMKII-activated sample, HeLa cells were treated with high [Ca^2+^] (1.26 mM), 10 μM ionomycin and 10 nM Calyculin A for 30 min to activate CaMKII. The CaMKII-inhibited sample was treated with 10 μM of KN-62 for 30 min, and subsequently treated with high [Ca^2+^] stimulation supplied with 10 μM of KN-62 for 30 min. Total lysates (1.5 mg) from each experimental condition was subjected to anti-KIF3A-immunoprecipitation overnight to pull down endogenous KIF3A, followed by SDS-PAGE and western blot analysis using anti-Phospho-serine/threonine antibodies to detect KIF3A phosphorylation status. 100 μg of total lysate was loaded as input. (D) HeLa cells were transfected with FLAG-KIF3A and FLAG-KIF3A-S689A respectively. The cells were treated similarly as 1(C) to activate or inhibit CaMKII. Total cell lysate (1 mg) was incubated with FLAG-M2 beads overnight to pull down the KIF3A constructs, followed by SDS-PAGE and western blot analysis using anti-Phospho-serine/threonine antibodies to detect KIF3A phosphorylation status. 100 μg of total lysate was loaded as input.

In this study, we use both the mouse and human KIF3A cDNA for our experiments. Hence, we mutated serine at position 690 to aspartic acid - S690D (and mouse S689D) to mimic the phosphorylated KIF3A and to alanine - S690A (mouse S689A) to mimic the non-phosphorylated KIF3A. In order to confirm the phosphorylation of CaMKII on serine-690/689 of KIF3A, we performed an *in vitro* kinase assay by incubating activated CaMKII with bacterial purified tail proteins of KIF3A wild-type (WT) or the phospho-mutants. FLAG-CaMKII protein was purified from HEK293 cells and activated by incubation with CaCl_2_, calmodulin (CaM) and ATP. The GST-KIF3A-tail of WT, S689A and S689D mutants were incubated with the activated CaMKII supplied with ATP. We observed that only the wild-type KIF3A-tail is highly phosphorylated by CaMKII, while both mutants were only weakly phosphorylated possibly at other serine or threonine residues within the C-terminal tail (Fig. 1B).

In order to confirm that KIF3A is phosphorylated by CaMKII *in vivo*, we incubated the cells in high [Ca^2+^] buffer to activate CaMKII and also used a small molecule inhibitor of CaMKII, KN62, to block kinase activity. Endogenous KIF3A was immunoprecipitated to ascertain its phosphorylation status. We found that KIF3A became phosphorylated in high [Ca^2+^] buffer and KN62 was effective in blocking this phosphorylation activity (Fig. 1C). We also found that unlike KIF3A-WT, KIF3A-S690A could not be phosphorylated in high [Ca^2+^] buffer suggesting that S690 is indeed the residue which is phosphorylated when the cells are incubated in high Ca^2+^ conditions (Fig. 1D). Taken together, our results suggest that CaMKII is the kinase responsible for phosphorylating KIF3A at serine-690 in the tail domain.

### Phosphorylation of KIF3A tail domain prevents motor-tail interactions

We next applied different biochemical and biophysical methods to determine if KIF3A adopts and autoinhibited state which can be relieved by phosphorylation. Bacterial purified His_6_-KIF3A-motor and the GST-tagged tail domains of mouse KIF3A-WT, KIF3A-S689A and KIF3A-S689D were mixed and subjected to GST-pull down. We found that the tail domain of wild-type and the phospho-dead mutant S689A bind well to the motor domain, while there is little or no interaction between the motor domain and S689D-Tail (Fig. 2A). Additional FLAG-immunoprecipitation (FLAG-IP) assays were performed to test interaction between KIF3A motor domain and the tail domains of KIF3A-WT, KIF3A-S689A and KIF3A-S689D. In this assay, FLAG-tagged KIF3A-motor construct was co-transfected with GST-tagged KIF3A-WT-Tail, KIF3A-S689A-Tail or KIF3A-S689D-Tail constructs into HEK293 cells. The cell lysates were then subjected to FLAG-IP and western analysis. The results showed that FLAG-KIF3A-motor can pull down wild-type and phospho-dead GST-S689A-Tail but not the phospho-mimic GST-S689D-Tail (Fig. 2B). These results showed that KIF3A motor domain interacts with wild-type and phospho-dead tail domains but not the phospho-mimic tail domain.

**Figure 2.**
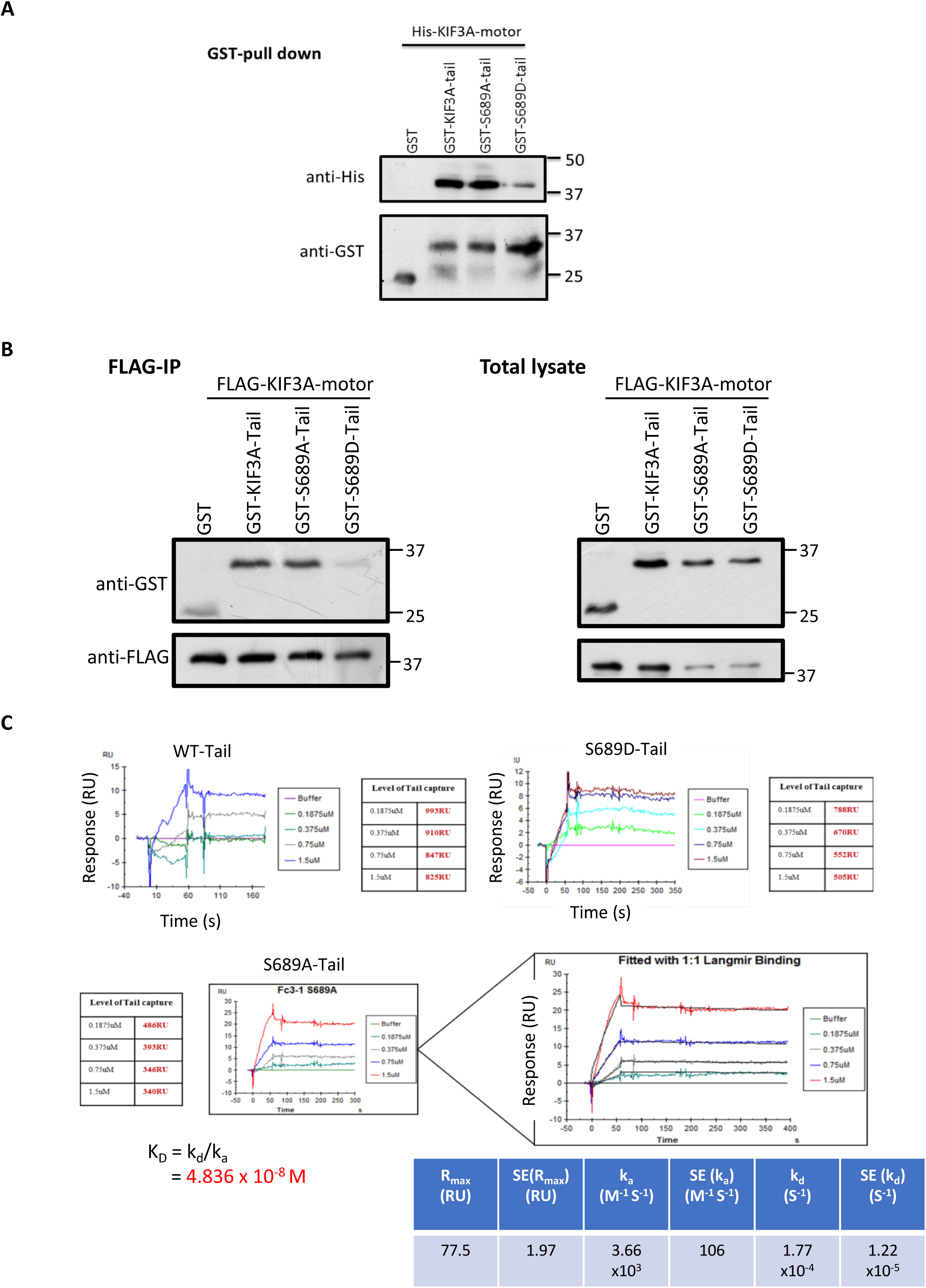
Interaction between the motor and tail domain of KIF3A. (A) Mixtures of bacterial purified His_6_-KIF3A-motor (5 μg) and 5 μg of purified GST-KIF3A WT-Tail, GST-S689A-Tail or GST-S689D-Tail were incubated with Glutathione Sepharose 4B. GST-fusion proteins were captured and the co-precipitated complex was separated by SDS-PAGE and analyzed by western analysis using anti-His and anti-GST antibodies. (B) FLAG-KIF3A-motor construct was co-transfected with GST, GST-KIF3A-Tail, GST-S689A-Tail and GST-S689D-Tail constructs in HEK293 cells. Total lysates were incubated with anti-FLAG M2 Affinity Gel and the eluents were detected by western blot using anti-FLAG or anti-GST. (C) SPR approach showed that the interaction between KIF3A-motor and S689A-Tail is stronger than WT-Tail, and S689D-Tail. GST-tail domains were captured on the chips by the immobilized GST antibody. Different concentrations of KIF3A motor were then injected into the Biacore system. GST-WT-Tail and GST-S689D-Tail show low binding intensity with KIF3A-motor. The sensograms are presented.

The binding affinities between KIF3A motor and tail domains in different phosphorylation states were further investigated by Surface Plasmon Resonance approach using the Biacore system. The interaction between KIF3A-motor and S689A-Tail was found to be stronger than that between the motor and tail domains of WT or S689D. The equilibrium dissociation constant, K_D_, of KIF3A-motor and S689A-Tail is estimated to be 4.836 x10^-8^ M, indicating a relatively strong binding. The binding signals of the motor domain with the WT- or S689D-Tail are too weak to calculate the K_D_ (Fig. 2C). From the binding profiles, we observed that the S689D tail domain showed the lowest affinity with the motor protein, while S689A showed the highest, suggesting that phosphorylation regulates the activity of KIF3A by disrupting its motor-tail interactions.

### Phosphorylation of KIF3A tail domain by CaMKII relieves KIF3A’s autoinhibition

Since S689A and S689D only show how KIF3A is regulated under simulated phospho-states, we wanted to observe how real phosphorylation regulates KIF3A. For this, we first carried out *in vitro* phosphorylation of the tail domain (Fig. 3A) and then checked the interaction between the motor and tail domains (Fig. 3B). Wild-type GST-KIF3A-Tail was first phosphorylated by purified CaMKII, then mixed with His_6_-KIF3A-motor and subjected to GST-pull down. We found that the motor protein can be pulled down by the non-phosphorylated tail domain but not by the tail domain that has been phosphorylated. When we increased the amount of the motor domain, there was still no interaction detected between the motor and phosphorylated tail domains (Fig. 3B). These results indicate that when KIF3A is not phosphorylated, it might adopt an autoinhibited state through motor-tail interactions and phosphorylation can relieve this autoinhibition.

**Figure 3.**
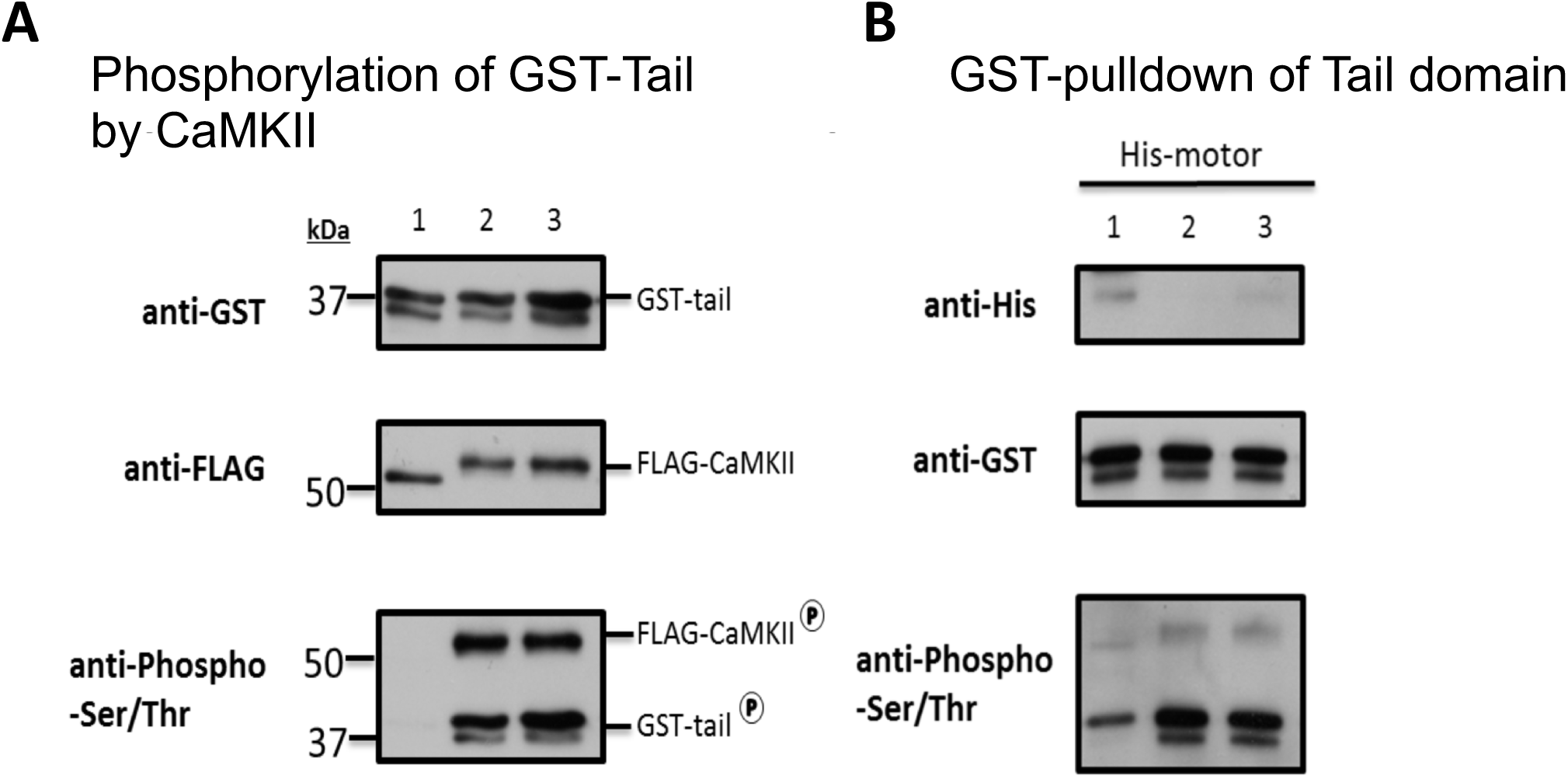
**In-vitro** kinase assay showed that phosphorylation inhibits KIF3A motor-tail interaction. (A) GST-WT-Tail domain phosphorylated by active CaMKII was detected by anti-phospho-Ser/Thr antibodies. Reaction mixtures were set up as follows: Lane 1: inactive FLAG-CaMKII + 5 μg GST-WT-Tail + 5 μg His-motor; lane 2: activated FLAG-CaMKII + 5 μg GST-WT-Tail + 5 μg His-motor; lane 3: activated FLAG-CaMKII + 5 μg GST-WT-Tail + 10 μg His-motor. (B) His-motor can be pulled down by the non-phosphorylated tail domain but cannot interact with the phosphorylated tail domain. The reaction mixtures were set up as in (A). The mixtures were subjected to GST-pull down, SDS-PAGE and western analysis using the respective antibodies.

### KIF3A goes through a conformational change when phosphorylated ***in vivo***

To study how the phosphorylation status affects autoinhibition of KIF3A in cells, a FRET (acceptor photobleaching) approach was adopted. FRET constructs with EYFP (enhanced yellow fluorescent protein) at the N-terminus and ECFP (enhanced cyan fluorescent protein) at the C-terminal of KIF3A were designed. The EYFP-ECFP construct acts as a positive control where the two fluorophores are close enough to cause FRET. EYFP-ΔTail-ECFP construct with the tail domain of KIF3A truncated acts as a negative control representing the extended KIF3A with no interaction between motor and tail domains. We propose that when KIF3A is in an extended state, EYFP and ECFP are too far away to cause FRET. When the motor and tail domains interact with each other, FRET could be detected (Fig. 4A).

**Figure 4.**
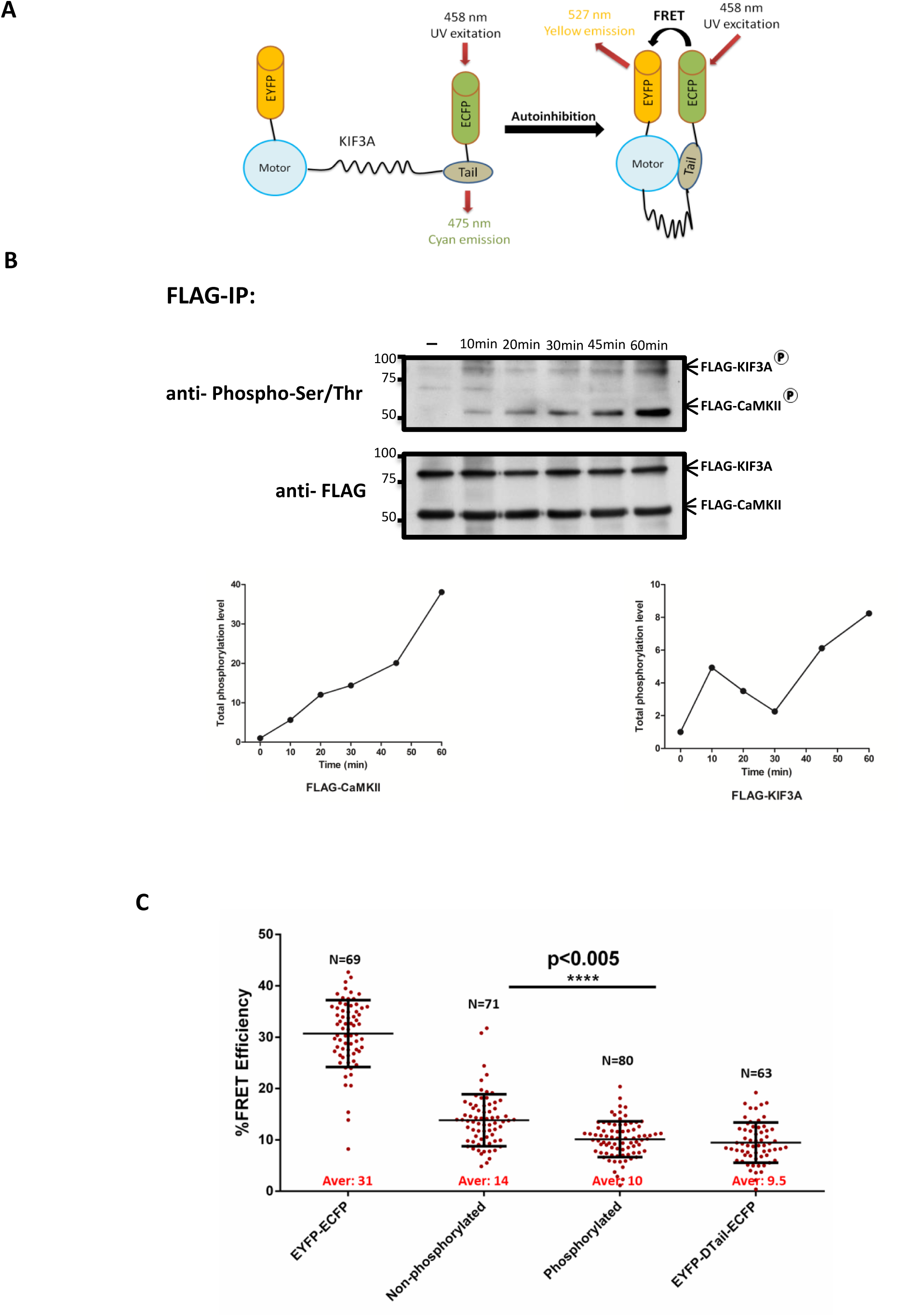
FRET efficiency of WT and mutants EYFP-KIF3A-ECFP. (A) Schematic diagram showing the KIF3A FRET construct. When KIF3A is in an autoinhibited state, EYFP and ECFP come close enough to cause FRET. (B) Phosphorylation levels of KIF3A and CaMKII at different time points. HeLa cells co-transfected with FLAG-KIF3A and FLAG-CaMKII were treated with high [Ca^2+^] buffer for 10, 20, 30, 45 and 60 min to activate CaMKII. The cells were then lysed and subjected to FLAG-IP. The phosphorylation levels of KIF3A and CaMKII were detected by anti-phospho-serine/threonine antibody. The graphs showing the band intensity of CaMKII and KIF3A at each time point were generated. (C) HeLa cells were transfected with *pEYFP-ECFP* as positive control, *pEYFP-ΔTail-ECFP* as negative control, and co-transfected with *pXJ-GST-CaMKII* and *pEYFP-KIF3A-ECFP*, which were further treated with low [Ca^2+^] buffer (non-phosphorylated KIF3A) and high [Ca^2+^] buffer (phosphorylated KIF3A). The graph shows the FRET efficiency of all samples. The number of cells analyzed (N) and statistical significance are indicated. The average FRET efficiency for each sample is in red.

In this experiment, HeLa cells expressing FLAG-CaMKII and FLAG-KIF3A were treated with high [Ca^2+^] buffer to stimulate the phosphorylation of CaMKII and KIF3A. In this set up, the phosphorylation levels of CaMKII and KIF3A in the cells kept increasing for 1 h after stimulation, suggesting a timeframe for carrying out the FRET experiment (Fig. 4B). To perform the acceptor photobleaching FRET experiment, the FRET construct EYFP-KIF3A-ECFP was co-transfected with GST-CaMKII into HeLa cells and acceptor photobleaching was performed using Zeiss 710 confocal microscope. At 24 h post-transfection, we treated the control cells (positive control) and experimental cells with extracellular Ca^2+^ to stimulate CaMKII, followed by the FRET experiment. We observed that under Ca^2+^/CaMKII stimulation, HeLa cells transfected with EYFP-KIF3A-ECFP exhibited lower FRET efficiency compared to the positive control and non-Ca^2+^ treated EYFP-KIF3A-ECFP transfected cells (Fig. 4C). Same experiment repeated in U2-OS cells also showed consistent results (Fig. S2A). To ensure that KIF3A is phosphorylated after Ca^2+^ stimulation, western blotting was done using cell lysates with co-transfected EYFP-KIF3A-ECFP in the presence of Ca^2+^, confirming phosphorylation of KIF3A was stimulated by high [Ca^2+^] buffer (Fig. S1). We observed a relatively low FRET efficiency for the non-Ca^2+^ treated EYFP-KIF3A-ECFP transfected cells, although the protein was expected to be non-phosphorylated and in a folded conformation. Nevertheless, the p-values between the non-phosphorylated and phosphorylated situations (p<0.005) are significant and indicate conformational change from the folded to extended states after phosphorylation. The average FRET efficiency of the cells co-transfected with KIF3A-motor-ECFP and EYFP-S689A-tail was also shown to be higher than cells transfected with wild-type or the phospho-mimic tails (Fig. S2B). These experiments suggest that KIF3A adopts an extended conformation in the cells when it is phosphorylated.

### ATP hydrolysis activity of KIF3A motor is inhibited by ADP-binding and ATPase inhibitors

Kinesins are ATPases that hydrolyze ATP to generate the energy required for stepping along microtubules. The motor domain of KIF3 catalyzes ATP hydrolysis. To determine whether the ATPase activity of KIF3A-motor is preserved in the recombinant protein, we first performed an ATPase assay using the recombinant KIF3A-motor domain. We found that recombinant KIF3A-motor protein was able to hydrolyze ATP efficiently. The hydrolysis activity was further increased after removing nucleotides from the motor protein. When the nucleotide-free motor protein was incubated with ADP again, there was a slight decrease in the ATP hydrolysis activity (Fig. 5A). The results suggest that the recombinant KIF3A-motor is enzymatically active and that trapped nucleotides lead to an inhibition of about 25%.

**Figure 5.**
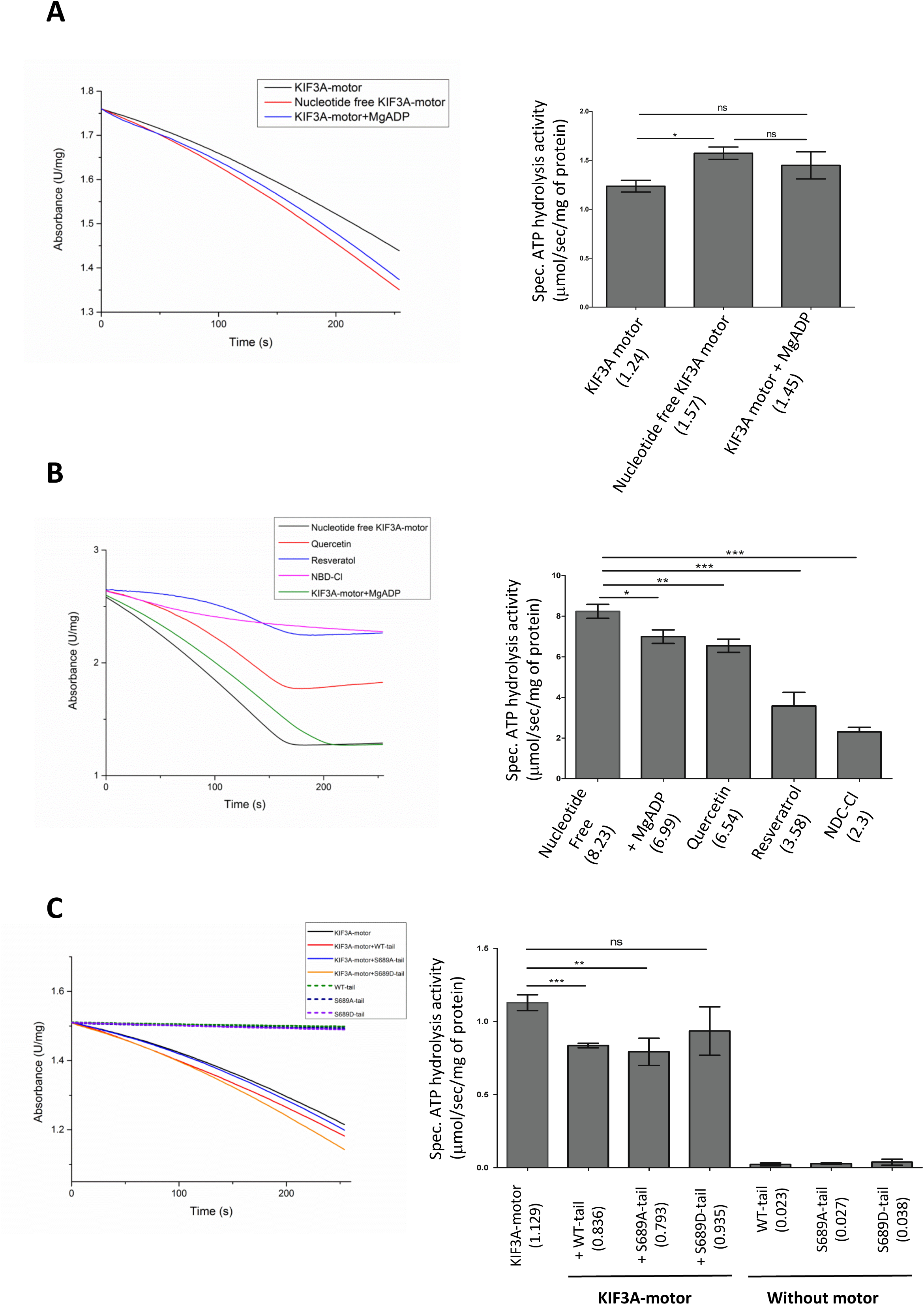
ATPase hydrolysis assay of the KIF3A-motor. (A) Left panel: the profile of UV absorbance of NADH at 340 nm (ATPase assays) of purified His_6_-KIF3A motor domain (black), nucleotide-free His_6_-KIF3A-motor domain (red) and His_6_-KIF3A-motor domain incubated with Mg^2+^-ADP (blue) over 5 min; right panel: the ATP hydrolysis activities of the three samples. (B) Left panel: the profile of UV absorbance of NADH at 340 nm (ATPase assay) of nucleotide-free His_6_-KIF3A-motor domain (black), His_6_- KIF3A-motor domain incubated with Mg^2+^-ADP (green) and His_6_-KIF3A-motor domain incubated with quercetin (red), resveratrol (blue) and NBD-Cl (pink) over 5 min; right panel: the ATP hydrolysis activity of the five samples. (C) Left panel: the profile of UV absorbance of NADH at 340 nm (ATPase assay) of nucleotide-free His_6_-KIF3A motor domain (black, line); His-KIF3A-motor domain incubated with GST-KIF3A-WT-tail (red, line), GST-KIF3A-S689A-tail (blue, line) and GST-KIF3A-S689D-Tail (orange, line); GST-KIF3A-WT-Tail (green, dash line), GST-KIF3A-S689A-Tail (blue, dash line) and GST-KIF3A-S689D-Tail (purple, dash line) over 5 min; right panel: the ATP hydrolysis activities of the five samples. The data in each graph are expressed as the mean of three replicates with error bars representing standard deviations. The statistical significances are indicated.

We also performed the ATPase assay to test the KIF3A-motor activity in the presence of known ATPase inhibitors. ATPase inhibitors quercetin, resveratrol or NBD-Cl were incubated with KIF3A-motor, and the ATPase activities were assayed. All three inhibitors reduced the KIF3A-motor activity, among which resveratrol and NBD-Cl showed more significant inhibition than quercetin (Fig. 5B). Previous studies have shown that these inhibitors inhibit ATP hydrolysis within the F_1_-domain of F-ATP synthases (27).

### Motor-tail interaction inhibits KIF3A-motor domain’s ATP hydrolysis activity

Previous studies have reported that a full-length *Drosophila* kinesin heavy chain exhibited a compact conformation through head and tail domain interactions, which led to inhibition of its ATPase activity (28). The ATPase activity of kinesin-1 was found to increase 2.4-fold after removing the C-terminal tail domain (29). In order to check whether KIF3A-motor-tail interaction affects KIF3A-motor’s activity, ATPase hydrolysis assays of motor-tail mixtures were performed. We measured the ATPase activities of the KIF3A motor by incubating the same amount of His_6_-KIF3A-motor with GST-KIF3A-Tail, GST-S689A-Tail or GST-S689D-Tail for 1 h. Reaction mixtures of GST-KIF3A-Tail, GST-S689A-Tail or GST-S689D-Tail alone were measured as control. We found that the GST-S689D-Tail slightly inhibited the motor’s activity, while the GST-KIF3A-Tail and the GST-S689A-Tail inhibited the motor activity significantly (Fig. 5C). Since we have shown that KIF3A motor interacts with wild-type or the S689A-Tail but not with the S689D-Tail (Fig. 2), our current observations from the ATPase assays suggest that motor-tail binding inhibits the motor’s ATPase activity. Taken together, our observations suggest that motor-tail interaction affects energy utilization of the motor domain.

### KIF3A tail peptide fits into the hydrophobic pocket of the motor domain dimer

Our experimental data strongly suggest that interaction between the motor and tail domains resulted in an auto-inhibited state for KIF3A. To further develop our proposed mechanism, we generated molecular models for KIF3A motor-tail interactions. Amongst all the kinesins that are thought to function via autoinhibition, Kinesin-1 is the only family member whose atomic structure of the motor domain dimer in complex with a peptide from its tail domain has been resolved at high resolution (PDB ID: 2Y65, 2.2Å) (14). This protein-peptide complex from *Drosophila melanogaster* provides critical insights into the mechanism of autoinhibition of Kinesin-1. The phosphorylation status of S689 (mouse) present in the tail domain of KIF3A modulates motor-tail interactions and consequently its activity. Sequence comparison of KIF3A homologs from different species shows that the equivalent region around S689 (mouse) is conserved (Fig. 6A). Significantly, we also observe that the ^690^AKP^692^ sequence motif is preserved in the tail peptide (^939^QAQAKPIRS^948^) of Kinesin-1 that interacts with the motor domain dimer. Based on these observations, we rationalize that the conserved tail peptide sequence ^686^RKRSAKPETV^695^ from KIF3A could potentially interact with the motor domain dimer in a manner similar to that observed in Kinesin-1. We accordingly used the Kinesin-1 structure as a template to model the motor dimer-tail peptide complex of KIF3A (Fig. 6B).

**Figure 6.**
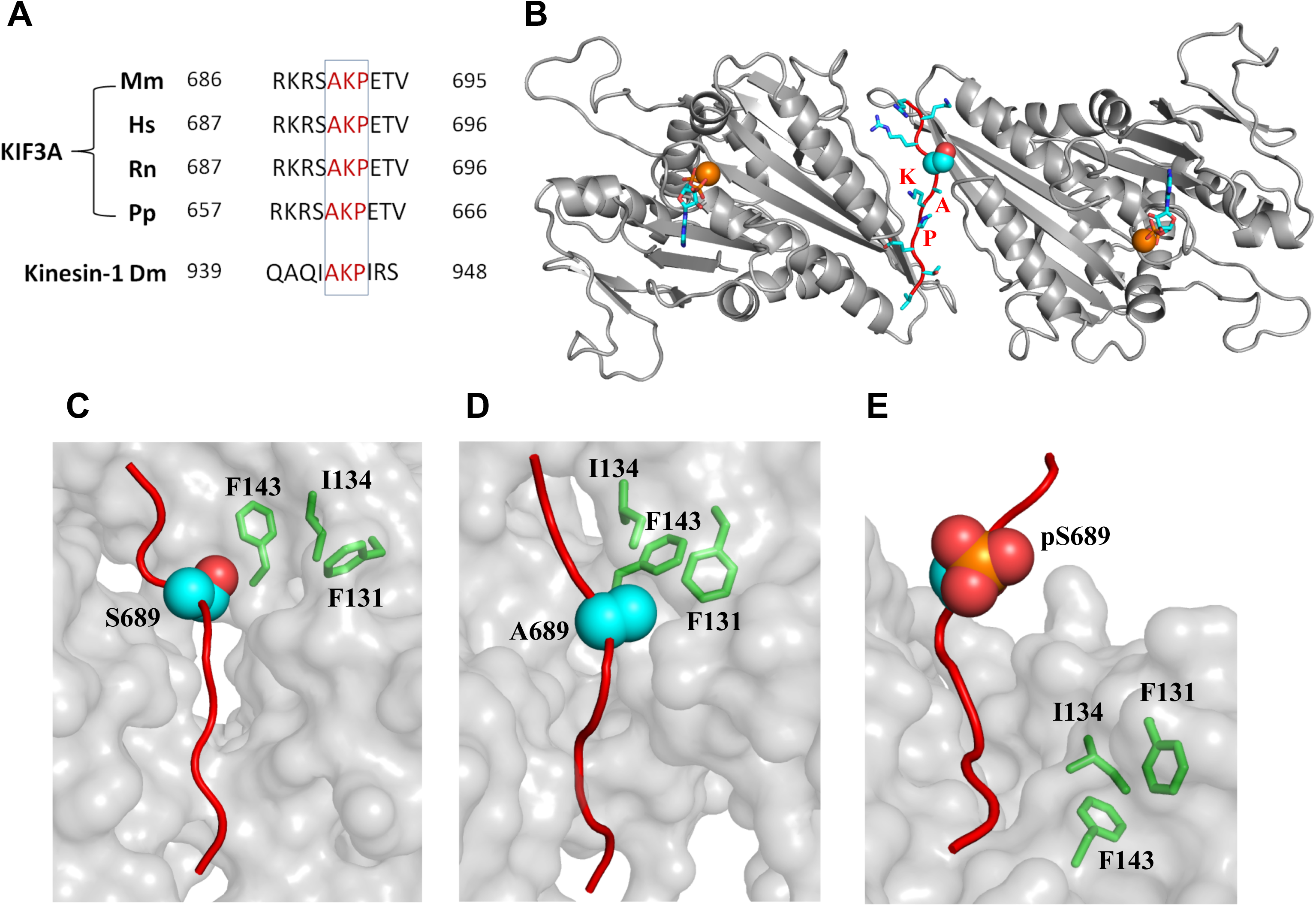
Molecular model of the motor-tail interaction of KIF3A. (A) Sequence alignment of KIF3A isoforms (Mm, *Mus Musculus;* Hs, *Homo sapiens;* Rn, *Rattus norvegicus;* Pp, *Poeciliopsis prolifica)* and Kinesin-1 (Dm, *Drosophila melanogaster)*. Amino acid sequence “AKP” are conserved between KIF3A and Kinesin-1. (B) Complex state molecular model of the mouse KIF3A peptide, residues 686-695 (shown in red cartoon backbone and cyan stick/sphere side-chain representation), and its motor dimer (gray cartoon representation) The “AKP” sequence is labelled and Serine-689 is specifically presented in sphere. ADP (cyan sticks) and Mg^2+^ ion (orange sphere), are also shown in the model. (C, D, E) Representative structure from molecular dynamic simulations showing the interaction between the motor domain dimer (gray surface representation) and the tail peptide (red cartoon) of WT (C), S689A (D) and phosphorylated S689 (E) state of the tail domains. Serine-689, Alanine-689 and serine-689 PO_3_^2-^ (pS689) is shown in sphere representation. The residues in the motor domain which form the hydrophobic pocket is shown in stick representation and labelled. Also see supplemental movies 1 to 3 for more details.

The tail peptide lies in the dimeric interface and localizes S689 into a hydrophobic pocket formed by residues F131, I134 and F143 on the motor domain (Fig. 6C). S689A mutation would enhance the local hydrophobic interaction between the tail peptide and the motor domain in this region whereas phosphorylation of S689 (pS689) would result in a not so favorable chemical environment due to the introduction of a negative charge in the vicinity of the hydrophobic pocket. This was further confirmed by MD simulations which showed that the peptide remained bound at the interface and the interaction between S689/A689 with the hydrophobic pocket was stable (Fig 6C, D; Supplemental Movies 1, 2). However, when a phosphate group (PO_3_^2-^) was added to S689 to mimic its phosphorylation, the stable interaction was broken primarily because of the introduction of the strong negative charge from the phosphate into the hydrophobic pocket (Fig 6E). Simulations show that the tail peptide does not remain stable and begins to disengage from the motor domain dimer interface (Supplemental Movie 3). These findings from the simulations support our observations that the KIF3A motor domain interacts with the tail domain in the dephosphorylated state of S689 whereas on phosphorylation of S689, this interaction is abrogated.

The Mg^2-^-ADP binding core is located at a site that is distant from the KIF3A motor-tail interacting interface (Fig. 6B), so the regulation of the ATPase activity by autoinhibition cannot be fully explained through this reduced model of the domain dimer-tail peptide complex. For Kinesin-1, it is proposed that the activity of the motor domain is regulated through a “double lockdown” mechanism (14). The cross-linking of the coiled-coil and the tail docking into the motor dimer interface together inhibit the motor’s MT-stimulated ATPase activity and its ADP release. For KIF3A, the regulation by the coiled-coil remains unknown. Since the phosphorylation state of S689 regulates the motor-tail interaction of KIF3A, a possible regulatory mechanism might be that the motor-tail interaction causes a local conformational change in the nucleotide binding pocket of the motor domain, triggering higher affinity towards ADP, resulting in blocking of ATP binding. Phosphorylation of S689 releases the KIF3A’s tail domain from the hydrophobic pocket of the motor domain, resulting in its activation.

## DISCUSSION

Kinesin’s activity has been reported to be regulated in many ways. They can be regulated through the control of gene expression or activation of kinesins during cell cycle by GTPase and kinases (30,31). One other regulatory mechanism that has been widely studied is autoinhibition (32). Kinesin in an autoinhibited state forms a folded, compact conformation that cannot hydrolyze ATP or associate with cargoes or microtubules. Activated kinesin which is released from autoinhibition transforms into an active, extended configuration. Kinesin-1, Kinesin-2 family members OSM-3 and KIF17, Kinesin-3 family member KIF1A and Kinesin-7 family member CENP-E have all been reported to adopt autoinhibited conformations by single-molecule analysis, biochemical assays, sedimentation assays or FRET (18,28,33-37). Currently there are two models, cargo binding and phosphorylation, that explain the activation mechanisms of autoinhibition. Kinesin-1 stays in a folded, inactive conformation in a cargo-free state. Cargo binding on the Kinesin-1 monomer relieves the autoinhibition and activates its MT gliding motility (38). Cargo binding activates KIF1A by driving the monomeric motors to form dimers, thereby activating its processive motility (39,40). For KIF11 and CENP-E, autoinhibition is reported to be regulated by phosphorylation. Phosphorylation of threonine-937 on KIF11 by CDK1 activates the motor and increases its MT-binding efficiency (17). Phosphorylation by TTK (dual specificity protein kinase) or CDK1-cyclin B causes CENP-E to unfold into an extended conformation and increase its processive motility along MT (18).

The phosphorylation status of serine-690 at the C-terminal region of human KIF3A has been shown to be regulated by the POPX2 and CaMKII phosphatase-kinase pair, and is shown to play a crucial role in KIF3A-mediated intracellular cargo transport (19). In this study, we showed that KIF3A-motor interacts with the wild-type and phospho-dead mutant tail domain, but not the phospho-mimic tail domain. Data from our FRET analysis also suggest that phosphorylation of serine-689 of mouse KIF3A tail domain triggers a conformational change from a folded configuration to an extended and active configuration in cells. We also found that motor-tail binding interferes with KIF3A’s ATPase activity. KIF3 motor complex is known to associate with its cargoes by the non-motor subunit KAP3 (41). Previous studies from our group have shown that the interaction between KAP3 and N-cadherin is not affected by phosphorylation. The phosphorylation status of serine-690 doesn’t affect the KIF3A-tail-KAP3 interaction either (19). In combination with our current results, we conclude that the phosphorylation state of KIF3A affects its ATPase hydrolysis activity rather than the cargo binding ability. Our previous study also showed that the *in vitro* KIF3A-microtubule binding was impaired when phosphorylation of serine-690 was precluded (19). Switch II loop/helix have been reported to interact with microtubules (42-45), which implies that the MT-binding site might also be located in the switch regions. The conformational change induced by phosphorylation might regulate the availability of both the ATP binding site and the MT-binding site. These findings suggest that phosphorylation may regulate both KIF3A’s MT-binding and force generation for MT-gliding. Molecular dynamics simulations of the conformational change of KIF3A’s tail peptide within the motor dimer’s interface showed that serine-689 or alanine-689 maintained high affinity with the hydrophobic pocket over simulation time, and the tail peptide stayed bound in the interface of the motor domain dimer (Figure 6 and Supplemental Movies 1, 2, 3). The addition of the phosphate group (PO_3_^2-^) to serine-689 destabilized the hydrophobic interaction, and the association between the tail peptide and the motor domain was no longer stable. The molecular mechanism suggested by the simulations support our observations that the KIF3A motor domain interacts with the tail domain, and phosphorylation of serine-689 releases the interaction. Based on the findings, we propose a model to explain how autoinhibition disrupts KIF3A-motor ATPase activity: in the inactive state, the tail domain folds into the motor domain dimer interface, and the motor and tail interact predominantly via hydrophobic forces. This interaction causes local conformational changes of the region involved in nucleotide binding, resulting in higher affinity towards ADP, thus inhibiting uptake of ATP. Phosphorylation of serine-689 in the tail domain breaks the stabilizing hydrophobic interactions between motor and tail. When the folded conformation is relieved, the catalytic core is once again free for ATP binding, resulting in the activation of KIF3A ATPase activity (Fig. 7).

**Figure 7.**
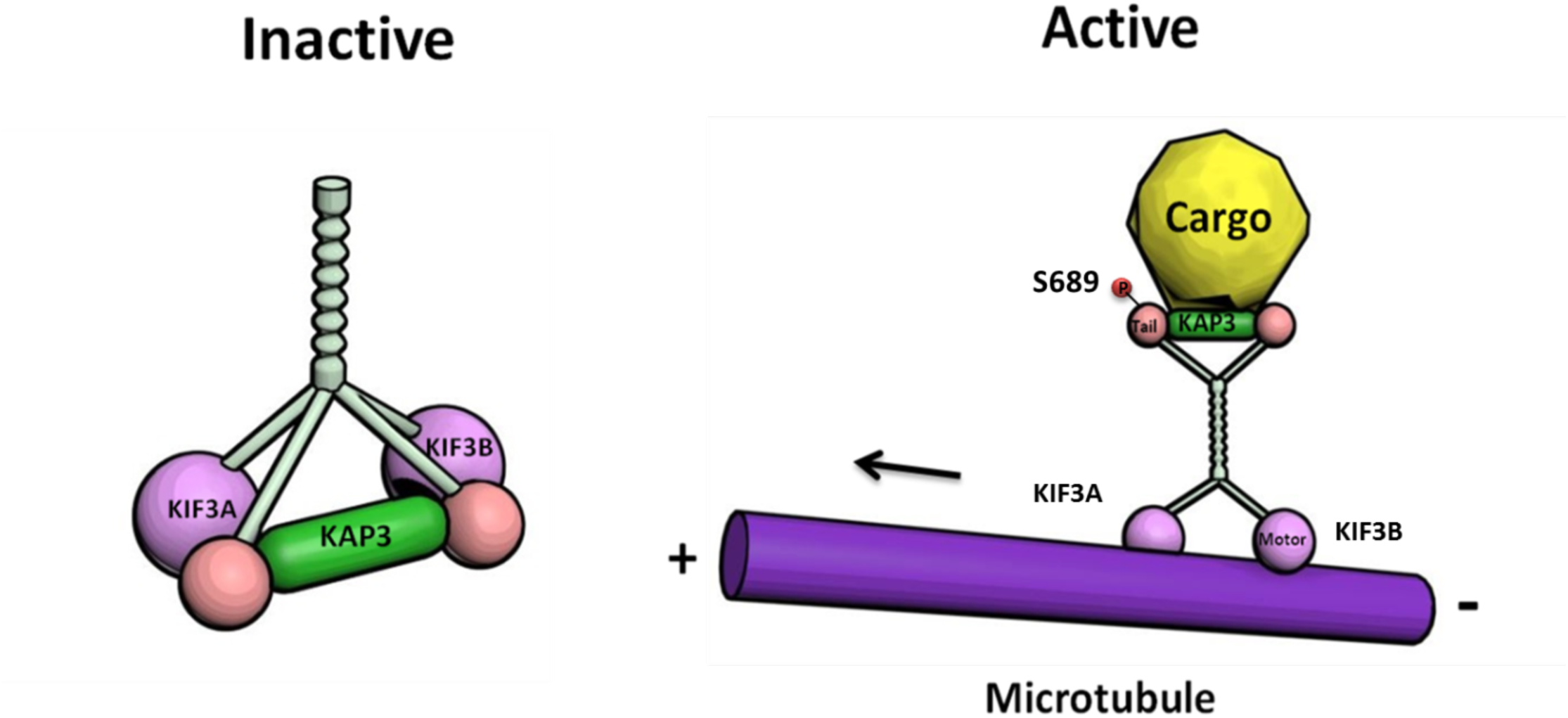
Proposed autoinhibition model of KIF3. When KIF3A is in an autoinhibited state, the motor and tail domains interact with each other and prevent KIF3A from binding to microtubules. Phosphorylation relieves KIF3A from autoinhibition. The active KIF3 motor can then bind to microtubules and transport cargoes along the microtubules.

The activity of the KIF3 complex is dependent on the coordination between KIF3A and KIF3B subunits. How the KIF3 complex is regulated in cells might be quite complex. There is no evidence to show whether KIF3B adopts an autoinhibited conformation. *Liang et al*. reported that phosphorylation of serine-663 in KIF3B disrupts its cargo-binding activity (46). KAP3 might also regulate KIF3’s activity. In addition to its cargo-binding role, some kinases or phosphatases regulate KIF3A or KIF3B via the interaction with KAP3 (47). Besides serine-689, there might be other phosphorylation sites that are essential for the regulation of KIF3A’s activity. There are still many questions to be answered for us to fully understand how the KIF3 motor complex is regulated. Further experiments are required to determine how the three subunits (KIF3A, KIF3B and KAP3), which make up the KIF3A complex, cooperate to allow proper functioning of the KIF3 motor.

## EXPERIMENTAL PROCEDURES

### Antibodies and plasmid constructs

Mouse monoclonal antibody to FLAG-M2 and rabbit polyclonal antibody to FLAG were purchased from Sigma-Aldrich. Mouse monoclonal antibody to GST was from Santa Cruz Biotechnology, and rabbit polyclonal antibody to GST was from Bethyl laboratories Inc (Montgomery, TX, USA). Rabbit polyclonal antibody to GFP was obtained from Abcam. Rabbit polyclonal antibody to His was from Cell Signaling Technology. Mouse monoclonal antibody to KIF3A was from BD biosciences. Rabbit polyclonal antibody against phospho-serine/threonine was from ECM Biosciences. *Mus musculus* full-length KIF3A (Addgene plasmid 13742) construct was purchased from Addgene (Cambridge, MA, USA). The motor domain of KIF3A (1-350) was amplified by PCR and cloned into the *pXJ40* vector with FLAG tag, or into a modified *pET30a-X* vector, yielding an N-terminal hexahistidine-tagged protein. The tail domain of KIF3A (598-701) was amplified by PCR and cloned into the *pXJ40* vector with GST tag, or was cloned into the *pGEX-6P-1* vector with an N-terminal GST tag. For the FRET constructs, the cDNA construct of EYFP was ligated in the *pECFP* vector with the ECFP tag. The full-length KIF3A (1-701) or the tail-truncated KIF3A (1-590) was amplified by PCR and ligated in *pEYFP-ECFP*.

### Cell culture and transfection

HeLa cells, U2-OS cells, COS7 cells and HEK293 cells were maintained in Dubecco’s modified Eagle’s Medium (DMEM) supplemented with 10% fetal bovine serum (FBS) at 37 °C in 5% CO_2_. Plasmid DNA were transiently transfected into cells reaching around 80% confluence using Lipofectamine™ 2000 (Invitrogen) according to the manufacturer’s protocols.

### Protein Expression and Purification

The plasmids *pET30aX-KIF3A-motor* and *pGEX-6P-1-KIF3A-tail* were transformed into *Escherichia coli* stain BL21 and grown in 1 liter of bacterial LB culture media with 500 mg/l Kanamycin at 37 °C till absorbance at 600 nm reached approximately 0.6. The bacterial culture was induced with 0.4 mM isopropyl β-D-thiogalactoside (IPTG) and incubated overnight at 160 rpm, 16 °C. The bacteria were pelleted by centrifugation at 4000 rpm for 40 min and stored in the −80 °C freezer.

For the purification of His_6_-tagged motor domain, harvested bacteria were suspended in protein lysis buffer containing 50 mM Tris-Cl pH7.5, 200 mM NaCl, 10% glycerol, DNase, RNase, supplied with 0.8 mM DTT and 2 mM Pefabloc and sonicated (25-30% power, 1 min × 3 times, with 2 min interruption). Clear lysate was loaded to Ni Sepharose™ 6 Fast Flow (GE Healthcare) bead slurry packed in an Econo-Pac^®^ Chromatography Column (Bio-Rad) and mixed for 1 hour. The resin was then washed with protein lysis buffer containing 20 mM imidazole. The proteins were eluted on a gradient from 50 to 500 mM imidazole. The eluted proteins were then concentrated and injected into HiLoad 16/600 Superdex 200 column (GE Healthcare). Proteins were eluted out by buffer containing 50 mM Tris-Cl pH7.5, 200 mM NaCl, 10% glycerol, 10 mM DTT, 2 mM MgCl_2_, and 2 mM CaCl_2_.

For the purification of the GST-tagged tail domain, harvested bacteria were suspended in protein lysis buffer containing 50 mM Tris-Cl pH7.5, 200 mM NaCl, 1.5% Sarkosyl, 1 mM lysosome, supplied with 1 mM DTT and 1 mM PMSF and sonicated (25-30% power, 1 min x 3 times, with 2 min interruption). Clear lysate was loaded to Glutathione Sepharose™ 4B (Amersham Biosciences) bead slurry packed in an Econo-Pac^®^ Chromatography Column (Bio-Rad) and mixed for 3 hours. The resin was then washed with protein lysis buffer. The proteins were eluted by elution buffer containing 20 mM reduced L-glutathione.

### Immunoprecipitation assay

Cells were co-transfected with *FLAG-KIF3A-Motor* and *GST-KIF3A-Tail* in 35 mm tissue culture dishes. The cells were collected 24 hours after transfection. Protein lysates were homogenized in lysis buffer containing 25 mM Tris-Cl pH7.4, 100 mM NaCl, 0.5% Triton X-100 and 1 x cOmplete Protease Inhibitor (Rhoche). The lysates were then clarified by centrifugation at 13,200 rpm for 10 min. 50 μl of ANTI-FLAG^®^ M2 Affinity Gel (Sigma) or Glutathione SepharoseTM 4B (Amersham Biosciences) bead slurry were added to the supernatant and mixed for 3 hours at 4 °C with constant rotation. Beads were then washed four times with washing buffer containing 25 mM Tris-Cl pH7.4, 500 mM NaCl. The bound FLAG-tagged or GST-tagged proteins were eluted by SDS sample buffer by heating at 100 °C for 10 min and subjected to SDS-PAGE and western blot analyses.

### Surface Plasmon Resonance (SPR)

Biacore 3000 (GE healthcare) was used to perform the SPR experiments. Flow cell (Fc) 1-4 surfaces were activated by injecting a mixture of 1-Ethyl-3-(3-dimethylaminopropyl)carbodiimide(EDC)/N-hydroxysuccinimide (NHS) (1:1) for 7 min at the flow rate of 10 μl/min. 30 μg/ml Anti-GST antibody in 10 mM NaOAc, pH 5, was immobilized to the chips for 4 min. Run ethanolamine through the chips to deactivate the remaining active esters for 7 min. Inject GST-KIF3A-Tail, GST-S689A-Tail and GST-S689D-Tail to Fc 2-4 for 3 min respectively, where Fc 1 acted as a reference. Following the capture of the tail proteins, recombinant GST was then injected at a flow-rate of 10 μl/min to block any remaining anti-GST binding sides to prevent non-specific interactions. Concentration series of 3 μM - 0.1875 μM His_6_-KIF3A-Motor protein was injected over the chip surfaces with the captured tail proteins. The surfaces were regenerated with 20 μl Glycine-HCl pH 2.1 and 10 μl 10 mM NaOH. The experiments were performed at 4 °C to keep the motor domain stable.

### In Vitro Kinase Assay

FLAG-CaMKII was expressed in HEK293 cells and subjected to FLAG-immunoprecipitation. FLAG-CaMKII was subsequently eluted out by 100 μg/ml 3X Flag peptide. Autophosphorylation of CaMKII was performed by incubating the eluted FLAG-CaMKII for 10 min at 30 °C with 1.2 μM calmodulin, 2 mM CaCl_2_ and kinase buffer containing 50 mM Tris-HCl, 10 mM MgCl_2_, 0.1 mM EDTA, 2 mM DTT, 0.01% Brij 35, pH7.5, supplied with 200 μM ATP. Purified GST-KIF3A-tail was phosphorylated by incubating with autophosphorylated CaMKII supplied with 200 μM ATP for 30-45 min at 30 °C with constant shaking. The reaction mixtures were subjected to SDS-PAGE and western blot, and the phosphorylation level was checked by anti-Phosphoserine/threonine antibody (1:500).

### Forster Resonance Energy Transfer

Plasmids pXJ-GST-CaMKII and pEYFP-KIF3A-ECFP were co-transfected into U2-OS or HeLa cells in two 35 mm dishes. One of the dishes with 50 μM [Ca^2+^] (140 mM NaCl, 5 mM KCl, 10 mM glucose, 20 mM HEPES, 2 mM EGTA, 1.13 mM MgSO4, 0.74 mM CaCl_2_, pH 7.36) was treated as control. To stimulate CaMKII phosphorylation, the other dish was treated with 50 μM [Ca^2+^] for 10 min and subsequently treated with 1.26 mM [Ca^2+^] (140 mM NaCl, 5 mM KCl, 10 mM glucose, 20 mM HEPES, 2 mM EGTA, 1.13 mM MgSO4, 4.38 mM CaCl_2_, pH7.36) supplied with 10 μM ionomycin and 10 nM calyculin. FRET experiments on the control and the phosphorylated samples were performed immediately after the treatments. Zeiss LSM 710 Confocal Microscope (Carl Zeiss) was used to perform the acceptor photobleaching FRET experiment. EYFP was excited by the laser at the wavelength of 514 nm with the emission filter BP of 525-551 nm. The gain was adjusted to make sure both channels reach saturation. ECFP was excited by the laser at 458 nm with an emission filter of 469-501 nm. A whole cell area was selected as the region of interest (ROI). Iteration was set at 120 and the sample was scanned for 10 cycles. The unbleached samples were used as control by scanning the cells for 10 cycles without bleaching. The photobleaching of EYFP within the ROI was achieved by scanning the region 10 times using 514 argon laser line. For the bleached samples, photobleaching was applied at the 6th scan cycle. The signals of ECFP within the ROI before and after EYFP bleaching were acquired. The intensity decrease of the EYFP channel shows the bleaching efficiency; the intensity increase of the ECFP channel indicates the FRET efficiency. FRET efficiency is determined by the formula,

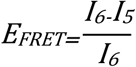

where I_5_ and I_6_ are the ECFP intensity of 5th and 6th time points, respectively.

### NADH-coupled ATPase Assay

10 μg of the purified KIF3A-motor domain in combination with the inhibitors or tail domains were added into 1 ml of the reaction solution containing ATP, phosphoenolpyruvate, pyruvate kinase, L-lactic acid dehydrogenase and NADH. NADH absorbance at 340 nm is recorded by Ultrospec™ 2100 pro UV/Visible Spectrophotometer (GE healthcare). The decline of NADH absorbance is used to monitor the rate of ATP hydrolysis. Data were normalized and the ATPase activity was calculated using

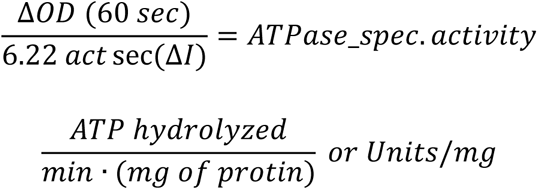

One unit corresponds to 1 μmol of ATP hydrolyzed/min.

### Modeling of KIF3A motor domain dimer: tail peptide complexes

The motor domain of mouse KIF3A (residues 1-350) was modeled via homology modeling techniques using the crystal structure of human KIF3B (PDB ID: 3B6U) as a template (Sequence Identity ~ 62%). The program MODELLER (version 9.13) (48) was used through the EasyModeller 4.0 graphical interface (49). The crystal structure of drosophila kinesin-1 motor domain dimer: tail peptide complex (PDB ID: 2Y65) was used as a template to construct a structural model for mouse KIF3A. For this, the modeled KIF3A motor domain was first superimposed onto the Kinesin-1 structure in order to generate the dimeric state of the motor domain of KIF3A. The peptide sequence “^686^RKRSAKPETV^695^” from the tail domain of KIF3A was then modeled by aligning the conserved “AKP” motif with the Kinesin-1 tail peptide “^939^QAQIAKPIRS^948^” present in the 2Y65 structure and subsequently changing the side-chains of flanking residues by in-silico mutations from Kinesin-1 amino acids to KIF3A amino acids. The motor domain dimer: tail peptide complex model of KIF3A thus obtained was refined by energy minimization. This optimized model was further used to generate two different complex variants with regard to the tail peptide sequence, one comprising of the phosphorylated state of S689 (pS689) and the other with the S689A mutation. Adenosine Diphosphate (ADP) and magnesium ion (Mg^2+^) present in the nucleotide-binding pocket of the motor domain in 2Y65 structure were preserved in all the three modeled complexes. All the structural superimpositions and *in-silico* mutations were performed using the Pymol Molecular Visualization Software (50).

### Molecular dynamics simulation of the complexes

The three different motor domain dimer: tail peptide complexes were subjected to molecular dynamics (MD) simulations using the PMEMD module and all-atom ff99SB force-field parameters (51) in the AMBER14 package. The AMBER library files for Phosphoserine and Adenosine Diphosphate (ADP) were downloaded from *http://research.bmh.manchester.ac.uk/bryce/amber* wherein the force-field parameters for the phosphorylated amino acid and the nucleotide were derived from the work of *Craft et. al*. (52) and *Meagher et. al*. (53) respectively. The parameter for magnesium ion (Mg^2+^) was used from the ions94 library of AMBER14. The N- and C-terminal ends of the polypeptide chains in the complex structures were capped with ACE and NME functional groups, respectively. The simulations were carried out in a cuboid box whose dimensions were defined by placing the complexes such that a minimum distance of 8 Å was maintained between any atom of the complex and box boundary. TIP3P water (54) was used for solvation and the net charge of all the systems was neutralized by adding appropriate number of chloride ions. Each of the three systems were energy minimized, heated to 300 K under NVT conditions, equilibrated for 500 ps and finally subjected to 100 ns of production dynamics in the NPT ensemble. Temperature (300 K) and pressure (1 atm) was regulated during the simulation using langevin dynamics (55) and weak coupling respectively (56). The PME method (57) was used to compute electrostatic interactions. Bonds involving hydrogen atoms were constrained with the SHAKE algorithm (58), enabling a time step of 2 fs for the time step for propagation of the dynamics.

### Statistical analysis

FRET and ATPase assay graphs and statistics were compiled using GraphPad Prism 5.01 and Origin 8.6. Other analyses were carried out using Excel software (Microsoft office 2010). All experiments were performed independently at least three times for statistical analysis. Significance was determined by two-way ANOVA for comparison between samples. *P* values < 0.05 were considered significant in all analyses.

## Data Availability

No datasets were generated or analysed during the current study.

## ACKNOWLEDGEMENT

We thank Dr Susana Geifman-Shochat and Mr Jediael Ng Zheng Ying (School of Biological Sciences, Nanyang Technological University) for their help with the SPR experiments. We also thank Dr. Chen Ming Wei and the Protein Purification Platform of NTU for providing help on protein purification. This work is supported by MOE Tier 1 (2014-T1-001-154 and 2016-T1-002-081). We thank A*STAR and NSCC for support.

## Author Contributions

K.C and C-G.K conceived the project. K.C, W.G.S and D.L. conducted the experiments. All authors analysed the results and reviewed the manuscript.

## Supplemental Information

**Supplemental Movies 1 to 3: Molecular dynamics simulation trajectories of tail peptide: motor domain complexes**. The motor domain dimer is represented in gray and the tail peptide in red ribbon representation. The side-chain of S689 (WT) (movie 1), A689 (S689A) (movie 2) and pS689 (phosphorylated S689) (movie 3) in the different states of the tail peptide is shown in stick representation. Adenosine di-phosphate (Stick representation) and Mg^2+^ (orange sphere) is also shown in the simulated trajectories. The WT and S689A state of peptide remains bound to the motor domain dimer whereas pS689 peptide dissociates from the complex.

**Figure S1.**
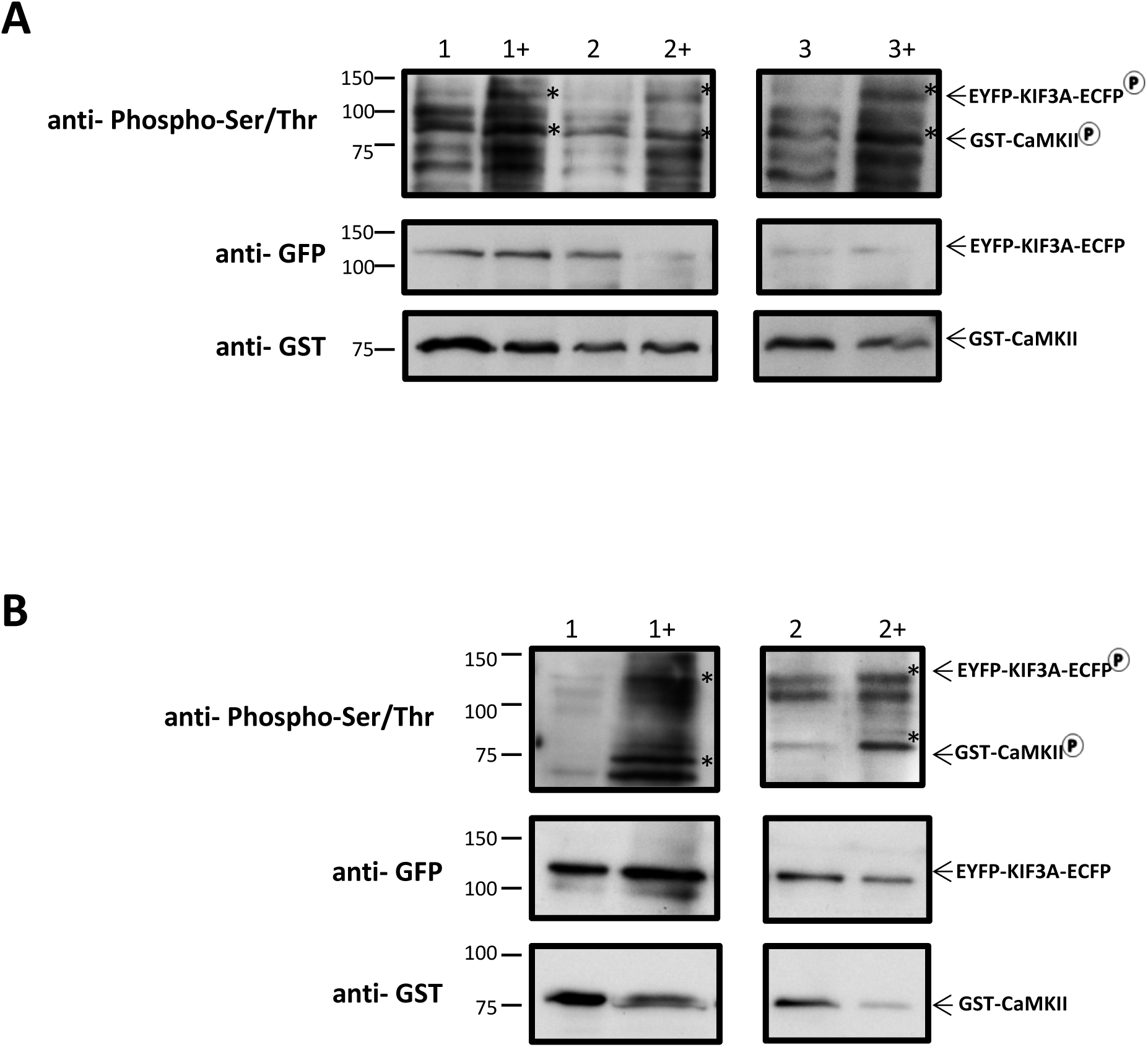
Phosphorylation of KIF3A after Ca^2+^ stimulation. U2-OS cells (A) or HeLa cells (B) were co-transfected with *pXJ-GST-CaMKII* and *pEYFP-KIF3A-ECFP*. Control or experimental cells treated with low or high [Ca^2+^] (indicated by “+”) buffers were lysed and sent for western-blotting (left). The phosphorylation levels of the total lysates were detected by phospho-serine/threonine antibody. The corresponding sizes of GST-CaMKII (80 kD) and EYFP-KIF3A-ECFP (125 kD) were indicated with asterix on the blots. Three independent experiments using U2-OS cells and two independent experiments using HeLa cells were performed. In all experiments, cells treated with high [Ca^2+^] buffer showed more intense phospho-KIF3A bands.

**Figure S2.**
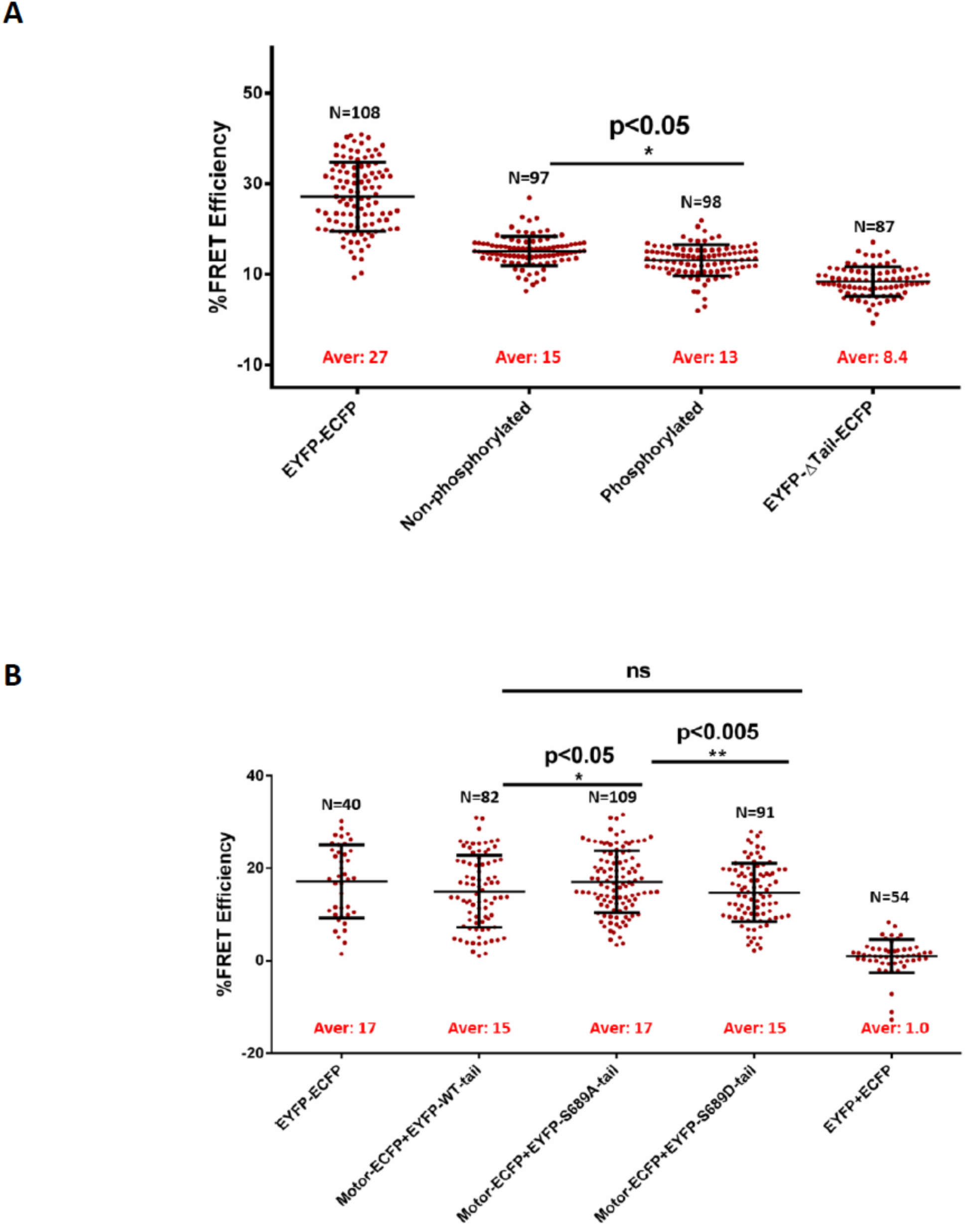
FRET efficiency of cells expressing KIF3A wild-type and mutants’ FRET constructs. (A) U2-OS cells were transfected with *pEYFP-ECFP* as positive control, *pEYFP-ΔTail-ECFP* as negative control, and co-transfected with *pXJ-GST-CaMKII* and *pEYFP-KIF3A-ECFP*. Control and experimental cells were treated with low or high [Ca^2+^] buffers. The graph shows the FRET efficiency of all samples was developed by combing the FRET efficiency of all the cells collected. The number of cells analyzed (N) and statistical significance are indicated. (B) FRET efficiency of cells expressing KIF3A-motor-ECFP and EYFP-tail of KIF3A WT and mutants. U2-OS cells were transfected with *pEYFP-ECFP* or co-transfected with *pEYFP* and *pECFP, pEYFP-KIF3A* and *motor-pECFP, pEYFP-S689A* and *motor-pECFP, pEYFP-S689D* and *motor-pECFP*, respectively. Three independent experiments were performed and the graph is developed by combing the FRET efficiency of all the cells collected. The number of cells analyzed (N) and statistical significance are indicated.

